# Alcohol induces mitochondrial fragmentation and stress responses to maintain normal muscle function in *Caenorhabditis elegans*

**DOI:** 10.1101/791814

**Authors:** Kelly H. Oh, Hongkyun Kim

## Abstract

Chronic excessive ethanol consumption produces distinct toxic and adverse effects on different tissues. In skeletal muscle ethanol causes alcoholic myopathy characterized by myofiber atrophy and loss of muscle strength. Alcoholic myopathy is more prevalent than all inherited muscle diseases combined. Current evidence indicates that ethanol directly impairs muscle organization and function. However, the underlying mechanism by which ethanol causes its toxicity to muscle is poorly understood. Here, we show that the nematode *C. elegans* recapitulates key aspects of alcoholic myopathy when exposed to ethanol. As in mammals, ethanol exposure impairs muscle strength and organization and induces the expression of protective genes, including oxidative stress response. In addition, ethanol exposure causes a fragmentation of mitochondrial networks aligned with myofibril lattices. This ethanol-induced mitochondrial fragmentation is dependent on mitochondrial fission factor DRP-1 (dynamin-like protein 1), and its receptor proteins on the mitochondrial outer membrane. Our data indicate that this fragmentation contributes to activation of mitochondrial unfolded protein response (UPR). We also found that robust perpetual mitochondrial UPR activation effectively counters muscle weakness caused by ethanol exposure. Our results strongly suggest that modulation of mitochondrial stress responses provides a mechanism to ameliorate alcohol toxicity and damage to muscle.

**Significance:** Chronic alcohol abuse causes the damage and toxicity to peripheral tissues, including muscle. Alcohol perturbs the structure and function of striated skeletal and cardiac muscles. These toxic effects of alcohol on striated muscles negatively impact morbidity and mortality to alcohol misusers. Here, we demonstrate that the nematode *C. elegans* also exhibits key features of alcoholic myopathy when exposed to ethanol. Ethanol exposure impairs muscle organization and strength, and induces the expression of genes that cope with alcohol toxicity. Particularly, we find that ethanol toxicity is centered on mitochondria, the power plants of the cell. As an adaptive protective response to mitochondrial dysfunction, ethanol-exposed cells induce global transcriptional reprogramming to restore normal mitochondrial function. Upregulation of this transcriptional reprogramming in *C. elegans* effectively blocks ethanol-induced muscle weakness, a key feature of alcoholic myopathy. Thus, the modulation of mitochondrial stress responses is a potentially promising therapeutic strategy to ameliorate alcohol toxicity to muscle.

## Introduction

Alcohol use disorder is a major health issue with enormous socio-economic costs. The detrimental effects of acute and chronic alcohol consumption on liver, pancreas, and brain are well documented (1). Chronic alcohol consumption also causes loss of skeletal muscle strength and cardiac contractile function (2, 3). Alcoholic myopathy and cardiomyopathy, collectively called alcoholic striated muscle disease, are more prevalent than all other inherited muscle disorders combined, with the incidence of 45-75% of chronic alcohol users (4). While chronic alcoholic myopathy can be reversed upon long-term abstinence from alcohol, alcoholic cardiomyopathy can be fatal as it becomes irreversible at a certain point and leads to heart failure. Despite such significant health impacts of alcoholic muscle disease, we do not fully understand the pathogenic mechanism by which alcohol exerts its toxicity on striated muscles at the molecular and cellular level.

Mitochondria are not only responsible for cellular energy production but also central to cellular signaling, metabolism, and cell death. These functions are closely linked to their architecture, which is dynamically regulated by fission and fusion processes in response to stress and metabolic state (5). Mitochondrial fission is mediated by the dynamin-related protein DRP1. DRP1 is recruited to the outer mitochondrial membrane via its receptor proteins, such as Fis1 and Mff, assembles into spirals to wrap around the circumference of mitochondria, and split the membranes into two through constriction (6) Mitochondrial fusion is also facilitated by dynamin family proteins. Whereas outer membrane fusion is mediated by the Mitofusins Mfn1 and Mfn2 (FZO-1 in *C. elegans*), inner membrane fusion is mediated by Opa1 (EAT-3 in *C. elegans*).

The mitochondrial unfolded protein response (UPR^mt^) is an adaptive transcriptional response elicited by various forms of mitochondrial dysfunction, such as a defect in mitochondrial protein import, an impairment of oxidative phosphorylation, and a perturbation of mitochondrial proteostasis (7). In *C. elegans* the bZIP transcription factor ATFS-1 (activating transcription factor associated with stress-1), a mammalian ATF-5 ortholog, is primarily responsible for mediating UPR^mt^ transcriptional response in the nucleus (8). Although ATFS-1 possesses both a mitochondrial targeting sequence (MTS) and a nuclear localization sequence, ATFS-1 is normally targeted to the mitochondria and undergoes degradation. However, when its mitochondrial import is reduced or blocked due to mitochondrial dysfunction, ATFS-1 is transported to the nucleus, where it directly activates the transcription of over 500 genes involved in mitochondrial quality control and cellular metabolism to restore mitochondrial proteostasis (9, 10). In addition to ATFS-1, the homeobox transcription factor DVE-1, along with its ubiquitin-like co-activator UBL-1, is also required for UPR^mt^ induction (11). Recent studies have shown that chromatin remodeling and epigenetic regulation are also critical for UPR^mt^ transcriptional regulation (12, 13).

*C. elegans* is well suited to study the genetic basis of alcoholic striated muscle disease. In addition to its genetic amenability, *C. elegans* body wall muscles are structurally analogous to mammalian striated muscles, and their structure and subcellular organelles can be easily observed through its transparent body in live animals. The mitochondria in body wall muscles are regularly organized along the myofiber easily discernable fashion and the mitochondrial fission and fusion machineries are conserved. Furthermore, cellular signaling, stress responses, and cell death, which are considered major components of alcohol toxicity pathways, are evolutionary conserved and well characterized.

In this report we show that *C. elegans* recapitulates key aspects of alcoholic myopathy; alcohol exposure causes mitochondrial fragmentation, induces oxidative stress and unfolded protein responses, and impairs muscle size and strength. Furthermore, our genetic analysis showed that the toxic effects of alcohol on *C. elegans* muscle result mainly from mitochondrial dysfunctions. Remarkably, we found that modulation of mitochondrial physiology by activation of mitochondrial unfolded protein response (UPR^mt^) protects from alcohol-mediated muscle dysfunction.

## Results

### Ethanol causes muscle dysfunction in *C. elegans*

To understand the toxic effects of ethanol on muscle function, we exposed *C. elegans* for 24 hours to a low dose of ethanol, which has a mild acute effect on locomotion, and then measured their movements in a liquid medium. While naïve control animals cultured in the absence of ethanol showed vigorous thrashing behavior, ethanol-exposed animals exhibited a significant reduction in the number of thrashing (Fig. 1A, Movie S1). This impaired thrashing does not result from neural depression caused by ethanol, since animals resistant to the intoxicating effects of ethanol still showed an impairment in thrashing (Fig. S1). The mobility of *C. elegans* in a natural habitat mainly consists of movements in the three dimensional space. To better simulate the natural locomotion, we performed a modified burrow assay (14), in which naïve control and ethanol-exposed animals are allowed to migrate across a solidified Pluronic F-127 gel layer from the bottom of a 12 well plate to an olfactory attractant on the surface. This assay has been shown to effectively detect even a slight deficit in neuromuscular function (14). We found that compared to naïve animals ethanol-exposed animals had difficulties in reaching the surface (Fig. 1B). In humans alcoholic skeletal myopathy leads to muscle fiber atrophy along with muscle weakness (3). We thus compared the sizes of striated body wall muscle cells in naïve and ethanol-exposed animals. Ethanol-exposed animals indeed had a smaller muscle than naïve animals (Fig. 1C and D). Furthermore, recovery after ethanol exposure on a regular culture plate for 4 hours did not restore the deficit in muscle size, indicating that muscle damage caused by ethanol exposure is not an acute temporary effect. This reduction in muscle size could be due to a reduced feeding. However, twenty four hour exposure of ethanol at this dose had no impact on the rate of pharyngeal pumping, indicating that a reduction of food intake by ethanol exposure is unlikely to cause a reduction of muscle function and size (Fig. 1 E).

**Fig. 1.**
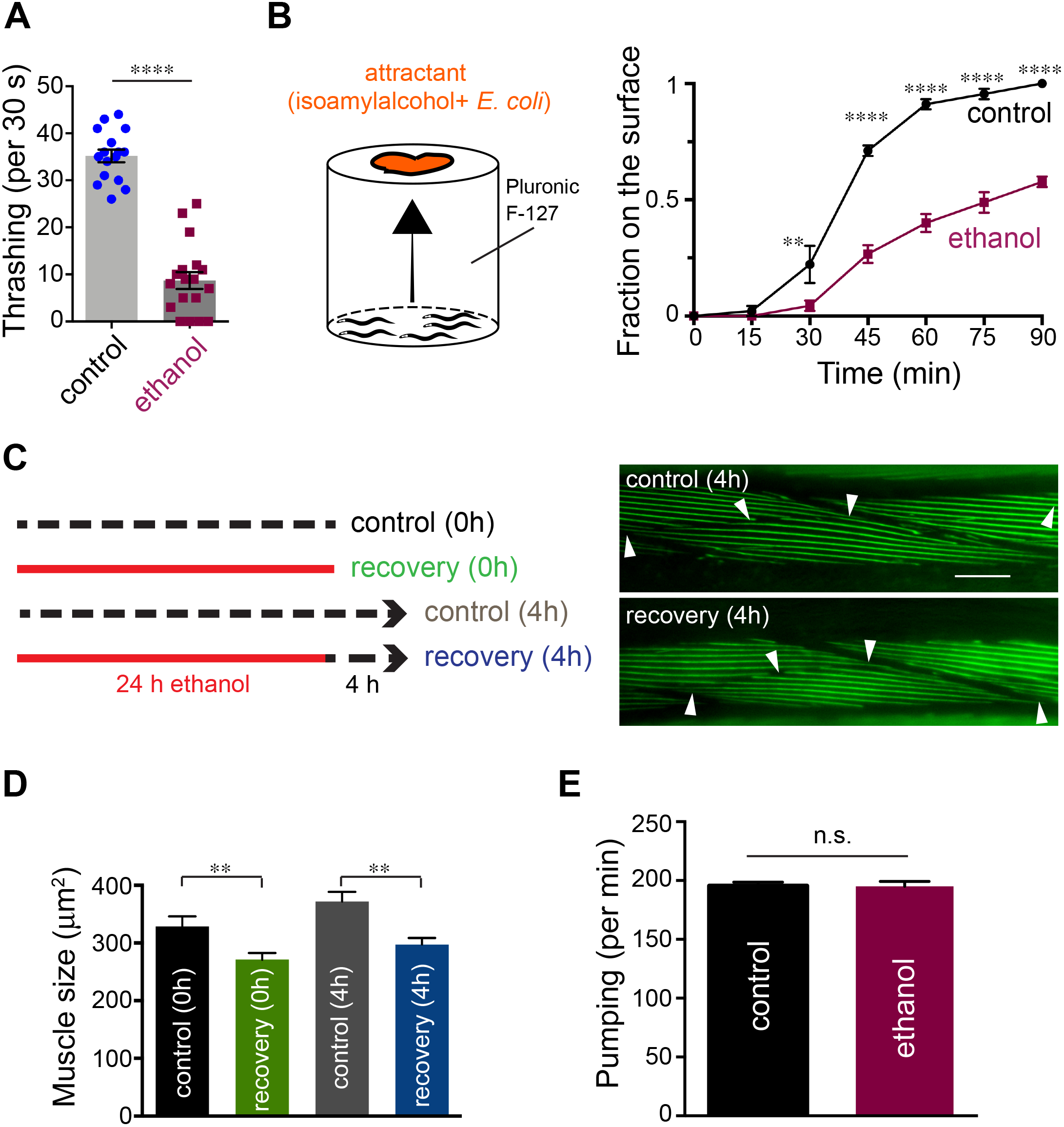
Long-term ethanol exposure causes muscle dysfunction. *(A)* Ethanol-exposed animals exhibit a reduction of thrashing frequency in a liquid environment, compared to control animals. ****p < 0.0001, unpaired t-test. *(B)* Ethanol-exposed animals exhibit a significant delay in burrowing across 26% Pluronic F-127 gel. Triplicates of 15 animals, **p < 0.005, ****p < 0.0001, two-way ANOVA, Sidak’s post-hoc analysis. *(C, D)* Ethanol-exposed animals persistently show smaller muscle size than their paired controls even after 4 hour recovery. Scale bar, 10 μm. Myosin fiber was visualized using transgenic animals expressing *myo-3*∷GFP (*cimIs46*). L4-stage animals (40 per group) were cultured in the absence (control 0h) or of presence (recovery 0h) of ethanol for 24 hours, and a half of them are recovered on NGM plates in the absence of ethanol for additional 4 hours. The borders of muscle cells are marked by arrowheads (mean ± SEM, t-test, *\*\*P* < 0.01). *(E)* Pharyngeal pumping rate remained the same in control and ethanol-exposed animals. Ten L4-stage animals were cultured in the absence (control) or presence of ethanol, and then their pharyngeal pumping rate was measured. p = 0.9, t-test

### Ethanol induces mitochondrial and ER stress responses in *C. elegans*

To explore how ethanol exposure causes the deficits in muscle function, we first determined which stress responses are induced in *C. elegans* after ethanol exposure using previously established reporter strains. All of the tested strains are listed in Materials and Methods. In these strains, the promoter sequences of a variety of stress response genes are fused to the fluorescent protein coding sequence, and the elevation of the fluorescent protein expression is an indicative of the induction of the stress response. As previously reported (15), robust induction of the alcohol dehydrogenase gene *sodh-1* reporter was observed (Fig. S2). We found that 24 hour exposure to ethanol increases the expression of the ER (*hsp-4p*∷GFP) and mitochondrial UPR (*hsp-6p*∷GFP) reporters, while not appreciably affecting the expression of the cytosolic stress reporter *hsp-16.2p*∷GFP (Fig. 2A). In addition, we observed that the expression of the oxidative stress reporter *gst-4p∷GFP* was induced by ethanol exposure (Fig. 2B). These results together indicate that ethanol induces genes that cope with alcohol detoxification, as in mammals.

**Fig. 2.**
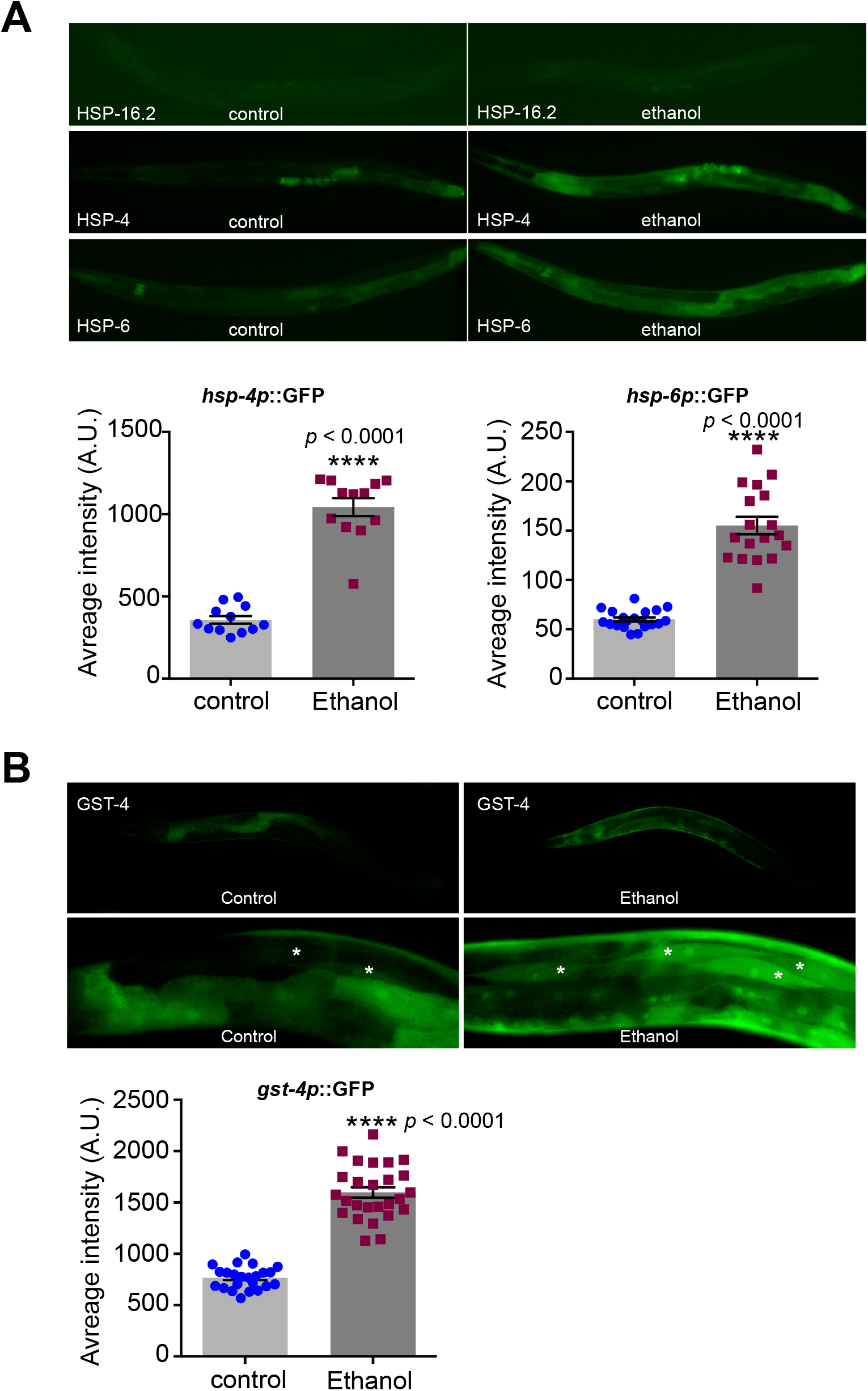
Chronic ethanol exposure induces stress response genes. *(A)* Mitochondrial and ER, but not cytosolic, stress responses are induced after 24 hour ethanol exposure. *hsp-6p*∷GFP, *hsp-4p*∷GFP, and *hsp-16.2p*∷GFP were used for reporters for UPR^mt^, UPR^ER^, and UPR^cyto^, respectively. In control animals, *hsp-6* and *hsp-4* are minimally expressed at some tissue. In ethanol-exposed animals, *hsp-6* and *hsp-4* are expressed throughout the entire body, including body wall muscle. mean ± SEM, t-test, ****P < 0.0001 *(B)* Ethanol exposure induces oxidative stress. *gst-4p*∷GFP was induced after 24 hour ethanol exposure. The white asterisks denote body-wall muscle cells. mean ± SEM, t-test, *\*\*\*\*P* < 0.0001

### Ethanol causes mitochondrial fragmentation in *C. elegans* muscles

Our observed stress response profile suggests that mitochondrial dysfunction might underlie ethanol-induced cellular stress, since mitochondrial dysfunction is known to cause oxidative stress and UPR^mt^ and indirectly UPR^ER^ (16, 17). Mitochondrial function is intimately linked to its dynamically regulated architecture; mitochondria undergo fusion and fission, and the balance of these two processes determines the mitochondrial architecture. Various cellular stressors alter mitochondrial function, and changes in mitochondrial structure enable the cell to restore the mitochondrial homeostasis (5). Therefore, we examined if ethanol exposure would alter the mitochondrial architecture in muscle.

To assess mitochondrial networks in striated body-wall muscle, we generated an integrated transgenic line that expresses a fusion protein at the outer mitochondrial membrane targeting sequence of TOMM-20 and FusionRed (*tomm-20*∷FusionRed) under the control of the muscle specific *myo-3* promoter. We observed that animals exposed to ethanol for 24 hours showed swollen mitochondria, fragmented mitochondrial networks, or complete disruption of mitochondrial networks in striated body-wall muscle (Fig. 3A). Mitochondrial fragmentation can result from either increased fission or decreased fusion. Therefore, we reasoned that if a reduction of fusion is responsible for ethanol-mediated mitochondrial fragmentation, significant mitochondrial fragmentation in fission defective animals is expected, and if increased fission is a major cause for the fragmentation, then fragmentation would be alleviated in fission defective animals. In *C. elegans* mitochondrial fission is mediated by DRP-1 and its receptors FIS-1/FIS-2 and MFF-1/MFF-2 (18, 19) (Fig. 3B). First, we tested whether muscle mitochondria undergo fragmentation in ethanol-exposed *drp-1(tm1108)* null mutant animals. Although it was not feasible to quantify the degree of fragmentation due to the highly connected mitochondrial network in *drp-1* mutant animals, ethanol did not change the mitochondrial architecture in *drp-1* mutant, demonstrating that DRP-1 is necessary for ethanol-induced mitochondrial fragmentation (Fig. 3C). Consistent with previous observations (18), naïve control *fis-1(tm1867)fis-2(gk414)* double mutant animals show a normal mitochondrial network structure. When *fis-1;fis-2* animals were cultured in the presence of ethanol for 24 hours, they show relatively mild mitochondrial fragmentation when compared to wild-type animals (Fig. S3A). We also examined *mff-1(tm2955);mff-2(tm3041)* double mutant animals for ethanol-mediated changes in mitochondrial architecture. Naïve control *mff-1;mff-2* mutant animals exhibit an increased interconnected mitochondrial networks (Fig. 3C and 3D), consistent with the fact that MFF proteins are a major receptor for DRP-1. Ethanol-exposed *mff-1;mff-2* mutant animals did not show any appreciable fragmentation and maintained interconnected mitochondrial networks similar to the control animals. Together, these results indicate that ethanol-induced mitochondrial fragmentation is primarily mediated by increased mitochondrial fission.

**Fig. 3.**
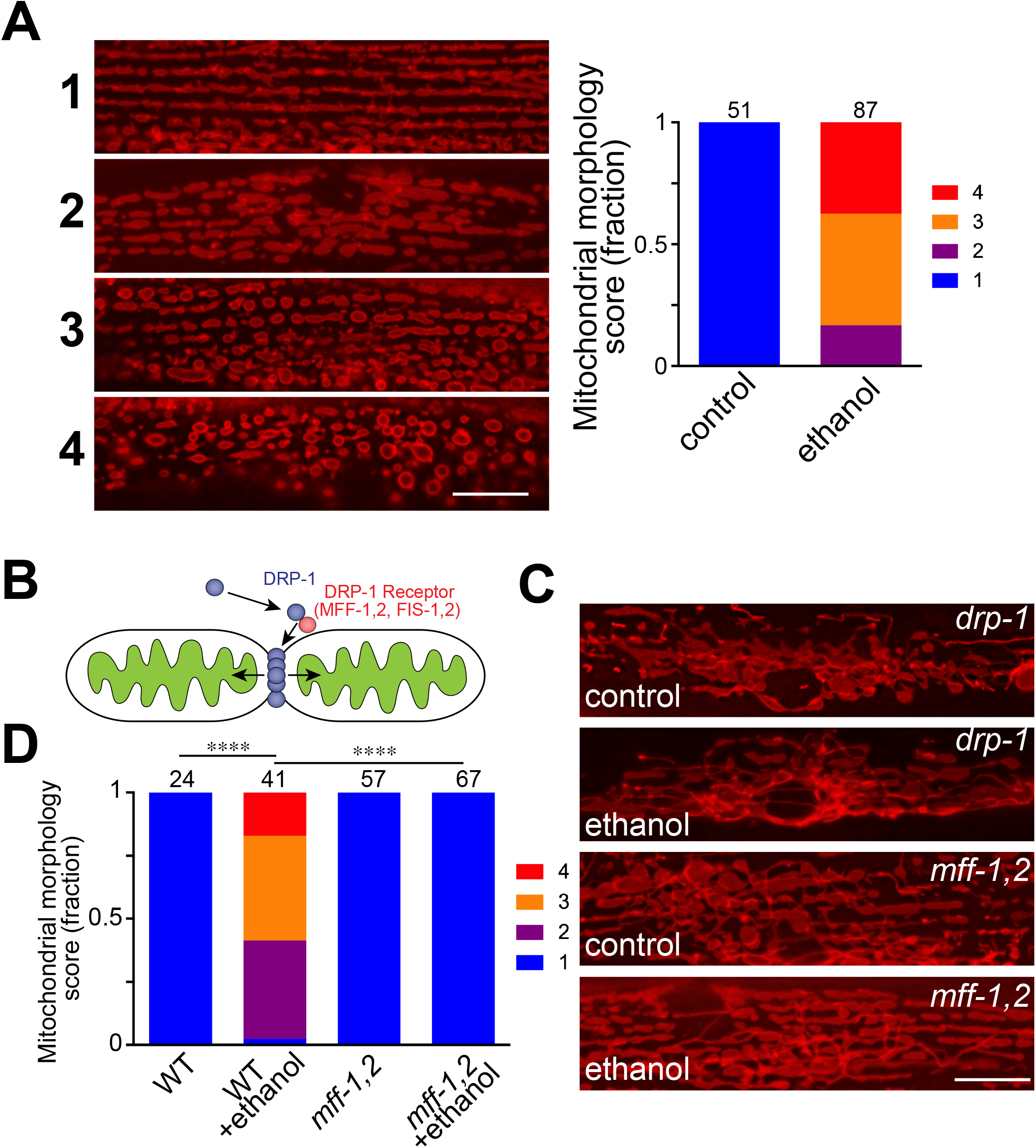
Ethanol-exposed animals exhibit muscle mitochondrial fragmentation in a fission machinery-dependent manner. *(A)* Muscle mitochondrial networks undergo fragmentation when *cimIs40[myo-3p∷tomm-20(1-68)∷*FusionRed] animals are exposed to ethanol for 24 hour. 1: normal linear networks, 2. moderate fragmented, 3. severely fragmentated but arranged in a linear fashion, 4. completely fragmented with no discernable linear position. Scale bar. 10 μm, p < 0.0001, Mann-Whitney test. The numbers above the bar graph are the sample numbers. *(B)* Schematic drawing of DRP-1 and DRP-1 receptors MFF-1,2 and FIS-1,2. *(C)* Mitochondrial networks in control and ethanol-exposed *drp-1* and *mff-1,2 (mff-1;mff-2)* double mutant animals. Scale bar, 10 μm. *(D)* Quantification of mitochondrial networks in wild-type and *mff-1,2* double mutant animals. ****p < 0.0001, Kruskal-Wallis test, Dunn’s post-hoc analysis. The numbers above the bar graph are the sample numbers.

We also determined whether mitophagy plays a role in ethanol-induced mitochondrial fragmentation (Fig. S3B). Mitochondrial fragmentation can be a process that separates damaged mitochondria from heathy mitochondrial networks for ultimate mitophagy (20). Thus, a deficit in mitophagy may block ethanol-induced mitochondrial fragmentation. However, even though the amount of mitochondria was considerably increased in these mitophagy mutant animals, mitochondria still underwent ethanol-induced fragmentation. Cell death mediators, such as CED-9 and CED-3, are known to influence mitochondrial fission and fusion (21, 22), but these mutant animals also exhibited alcohol-induced mitochondrial fragmentation comparable to wild-type counterparts (Fig. S3C). Together these results indicate that genes that mediate mitophagy and cell death are not involved in ethanol-induced mitochondrial fragmentation.

### Excessive mitochondrial fragmentation leads to the induction of UPR^mt^

Because long-term ethanol exposure in *C. elegans* causes both mitochondrial stress response and fragmentation, we explored whether each has any effect on the other. Therefore, we measured the levels of UPR^mt^ in *drp-1* and *fzo-1* mutant animals, which have defects in mitochondrial fission and fusion, respectively. Both *drp-1* and *fzo-1* mutant animals exhibited higher levels of UPR^mt^ reporter expression than the wild type control (Fig. 4A). These results show that perturbation of mitochondrial structure by a defect in either fusion or fission elicits mitochondrial stress. After ethanol exposure, the reporter expression was further increased in *drp-1* mutant animals that have highly connected mitochondria, but not in *fzo-1* mutants that have highly fragmented mitochondria (Fig. 4A). We also observed that the expression of the oxidative stress response gene reporter *gst-4p∷GFP* behaved similarly to the UPR^mt^ reporter (Fig. 4B). Together, these results suggest that disruption of normal mitochondrial networks causes UPR^mt^ and oxidative stress, and that mitochondrial fragmentation contributes to the ethanol-induced UPR^mt^ and oxidative stress response activation.

**Fig. 4.**
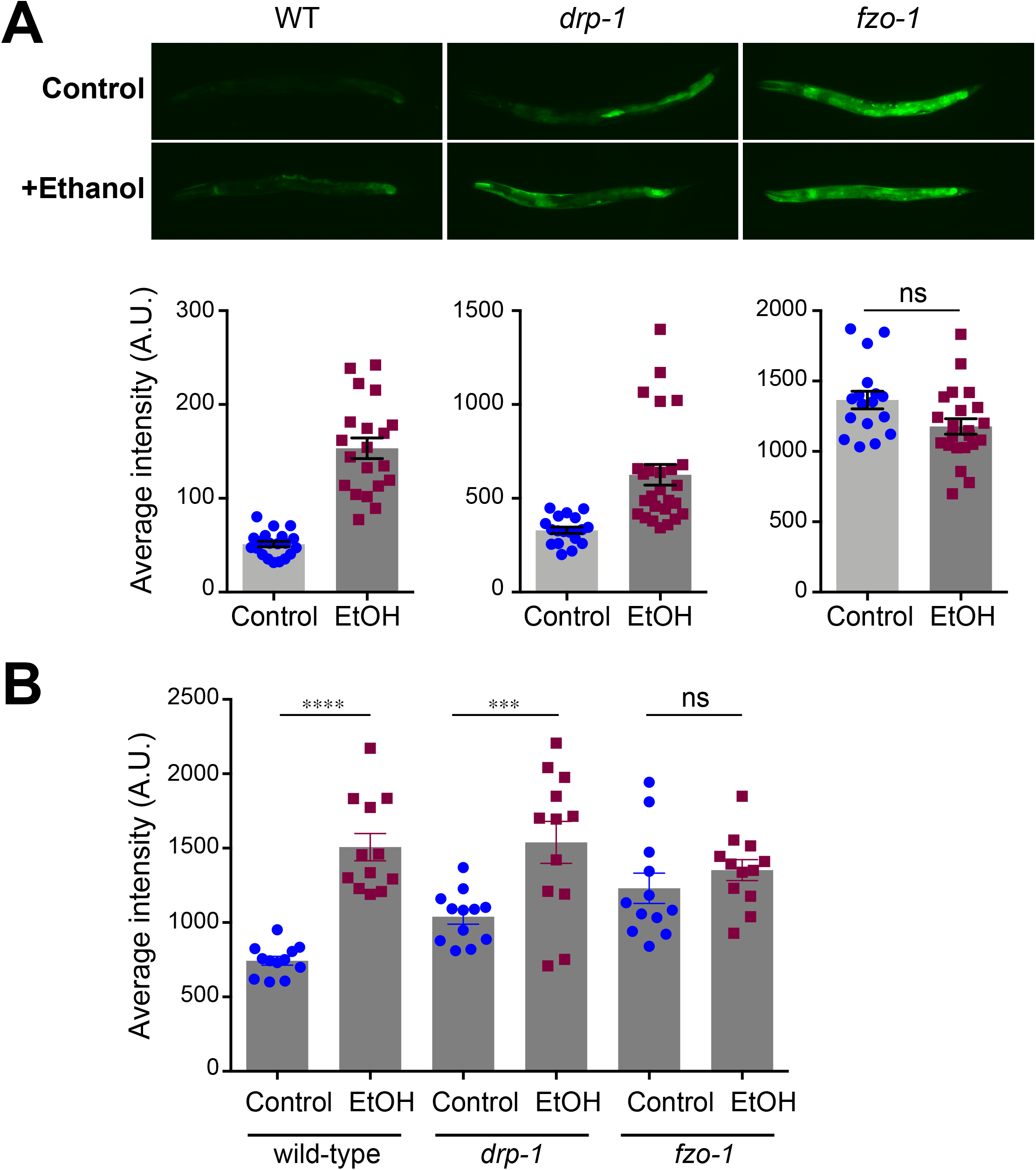
Alcohol-induced mitochondrial fragmentation contributes to UPR^mt^ and oxidative stress response. *(A)* UPR^mt^ (*hsp-6p*∷GFP) is up-regulated in both *drp-1* and *fzo-1* mutants, but UPR^mt^ in *fzo-1* mutant animals is not further increased by ethanol exposure. mean ± SEM ****p <0.0001, ***p<0.001, t-test. *(B)* Oxidative stress response (*gst-4p*∷GFP) is up-regulated by ethanol exposure in wild-type and *drp-1* mutant animals, but not in *fzo-1* mutant animals with hyper-fragmented mitochondria. mean ± SEM ****p<0.0001, ***p<0.0008, ns = 0.867, One-way ANOVA, Sidak’s pos-hoc analysis.

### Activated UPR^mt^ counteracts ethanol-induced muscle dysfunction

ATFS-1 is a master regulator of UPR^mt^ and its nuclear localization is an indicator of mitochondrial dysfunction (23). We determined whether ethanol exposure increases nuclear localization of ATFS-1 using a GFP-tagged ATFS-1 transgene. When these transgenic animals were cultured in the absence of ethanol, ATFS-1 was barely detectable due to an efficient degradation under normal condition (Fig. 5A), as previously reported (23). After 24 hour ethanol exposure ATFS-1 was found in the nuclei, albeit weak (Fig. 5A), similarly to a previous report (23). In addition to ATFS-1, we examined a co-regulator of UPR^mt^, DVE-1, which translocates into the nucleus upon mitochondrial stress (11), and found that ethanol exposure increases nuclear localization of DVE-1 (Fig. 5B). These results indicate that ethanol-induced UPR^mt^ is mediated by ATFS-1 and DVE-1.

**Fig. 5.**
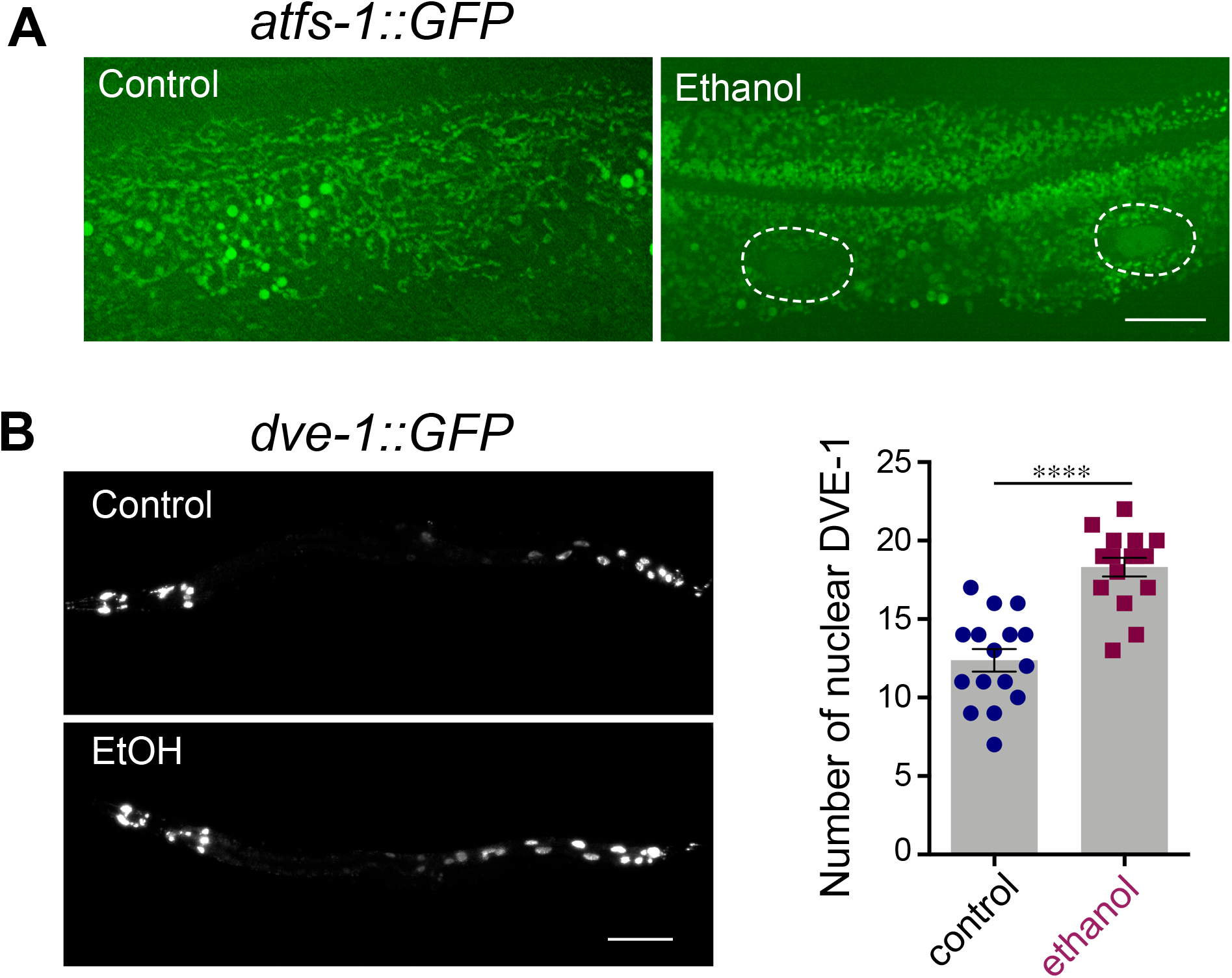
Ethanol activates transcription factors for UPR^mt^. *(A)* Ethanol exposure for 24 hour mobilizes ATFS-1∷GFP to the nuclei in the intestine. The dotted circles denote intestinal nuclei. *(B)* Nuclear DVE-1, which is required for the induction of *hsp-6* upon mitochondrial stress in addition to ATFS-1, is also increased upon ethanol exposure. Strain SJ4197 (*zcIs39[dve-1p∷dve-1∷gfp]*) was used. Number of DVE-1 positive nuclei was counted excluding head region, where DVE-1∷GFP is constitutively present in nuclei. mean ± SEM ****p<0.0001, t-test.

We hypothesized that the activation of UPR^mt^ is a homeotic mechanism to cope with ethanol toxicity and therefore UPR^mt^ is protective, at least short term, from ethanol toxicity. Thus, we compared muscle function of wild type, *atfs-1(lf, loss of function)*, and *atfs-1(et15gf, gain-of-function*) mutant animals using the thrashing and burrowing assays (Fig. 6, Movie S1). Under control condition, neither *atfs-1(lf)* or *atfs-1(et15gf*) mutant animals were significantly different from wild-type animals in thrashing frequency and burrowing ability (Fig. 6). After exposed to ethanol for 24 hours, both wild-type and *atfs-1(lf)* mutant animals showed diminished thrashing and burrowing. By contrast, ethanol-exposed *atfs-1(et15gf)* mutant animals showed strikingly robust thrashing and burrowing comparable to their naïve control counterparts, demonstrating that *atfs-1(et15gf)* mutant animals maintained muscle function after ethanol exposure (Fig. 6, Movie S1). Together these results indicate that perpetual UPR^mt^ activation protects animals from muscle weakness caused by ethanol toxicity.

**Fig. 6.**
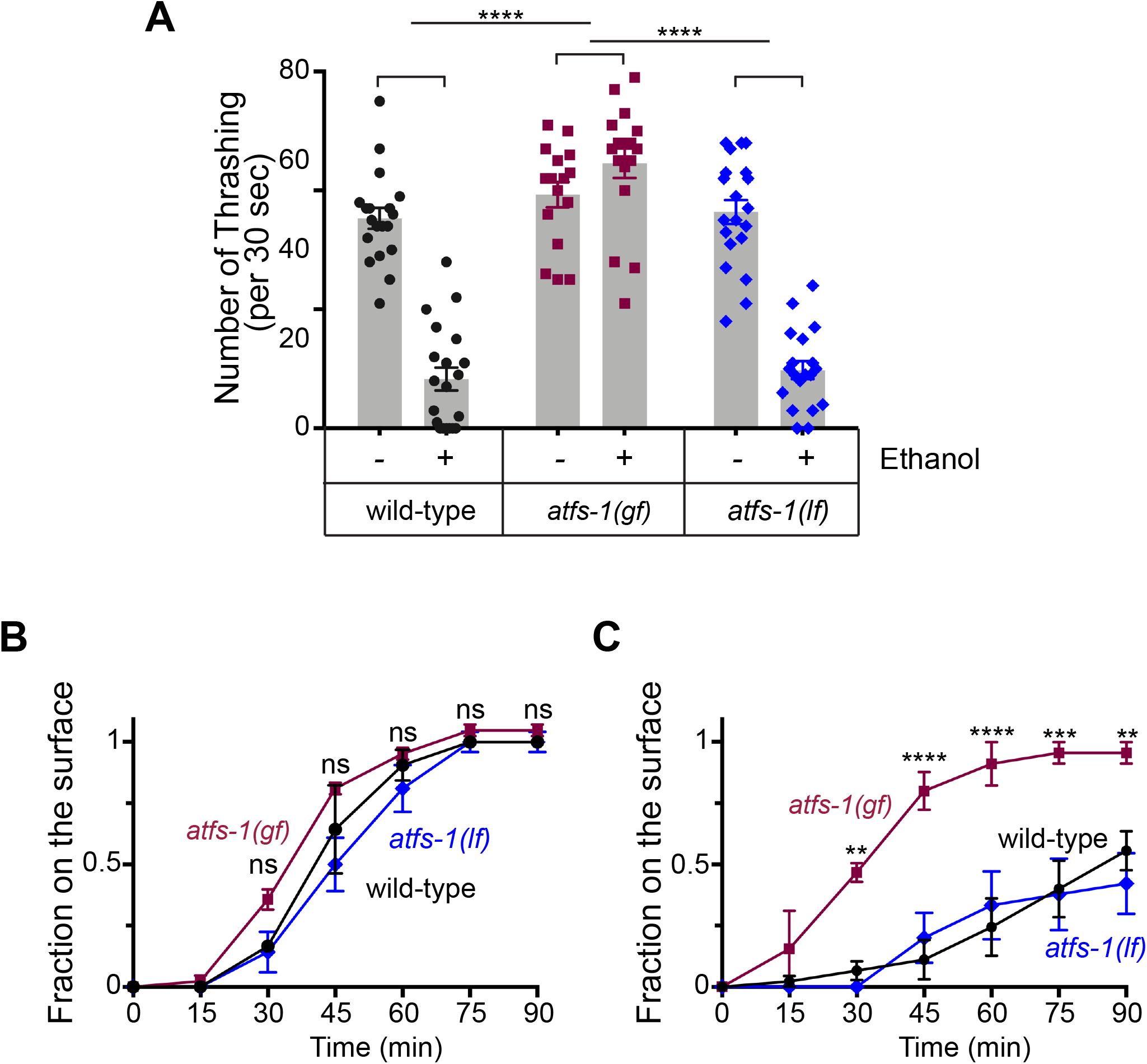
Constitutive activation of ATFS-1 protects from ethanol-mediated muscle weakness. *(A)* The thrashing assay post 24 hour ethanol exposure shows that *atfs-1(gf, gain-of-function)* mutant animals, but not wild-type or *atfs-1(lf loss-of-function)*, maintain normal thrashing. mean ± SEM ****p <0.0001, 2-way ANOVA, Tukey’s post-hoc analysis. *(B)* Age-matched wild-type, *atfs-1(gf)*, and *atfs-1(lf)* animals exhibit normal burrowing ability when grown under normal condition. *(C)* After 24 hour ethanol exposure *atfs-1(gf)* animals, but not wild-type or *atfs-1(lf)*, exhibit normal burrowing ability. Triplicate of 15 animals, mean ± SEM **p<0.01, ***p <0.001, ****p <0.0001, 2-way ANOVA, Tukey’s post-hoc analysis.

## Discussions

Chronic alcohol abuse leads to skeletal and cardio myopathy in humans. It is also well known that striated muscles of alcoholic patients show abnormalities in mitochondrial structures, defects in myofibril contractility, and metabolic reprogramming (3). In this study we employed two methods to measure muscle function, thrashing and burrowing assays. Unlike locomotion on a standard nematode culture dish, these two environments are more energetically challenging and therefore functional capacity of muscles can be readily examined. Using these two measurements, we demonstrate that ethanol-exposed animals recapitulate functional deficits in muscles as seen in alcohol-induced skeletal myopathy.

It has been noted that stress response pathways are activated in alcoholic myopathy (24, 25). The availability of various reporter strains to probe the activation of a specific stress response allowed us to demonstrate that ethanol exposure induces mitochondrial stress, ER stress, and oxidative responses. This response profile suggests that the primary target of ethanol may be the mitochondria. It is known that mitochondrial stress can lead to ER stress (16, 17) and mitochondria are a major player in oxidative stress pathway. Mitochondrial dysfunction can also explain the functional deficit of muscles since mitochondria are the primary source of energy production.

Mitochondrial architecture is closely linked to its function; highly connected mitochondria are more efficient energy producers and fragmented mitochondria are inefficient at oxidative phosphorylation. We discovered that ethanol exposure leads to mitochondrial fragmentation in *C. elegans*. Such ethanol-induced mitochondrial fragmentation was observed in human and rodent cardiomyocytes and skeletal muscles (26–29). The mitochondrial fragmentation can be the result of excessive fission or deficient fusion of mitochondria. Mutations in fission factors *mff-1/2* inhibited the ethanol induced mitochondrial fragmentation efficiently, while mutations in *fis-1/2* led to partial inhibition. FIS-1/2 has been known as fission factors, but recently they have been shown to increase mitochondrial fragmentation by inhibiting fusion rather than activating fission process directly (30). This may explain our observation that *fis-1/2* mutation was less effective in preventing ethanol induced mitochondrial fragmentation than *mff-1/2* mutation, which is likely to facilitate the fission directly. This further supports the notion that increased fission contributes to ethanol-induced mitochondrial fragmentation more than reduction of fusion, although we cannot completely rule out the possibility that mitochondrial fusion is also affected by ethanol.

We observed that the expression of *hsp-6p*∷GFP was elevated in both *drp-1* and *fzo-1* mutant animals, but the expression level was further increased by ethanol exposure only in *drp-1* mutant animals. Although we and others have shown that the outer mitochondrial membrane of *drp-1* mutant animals is highly connected, it was reported that the mitochondrial matrix appears to be compartmentalized (31), suggesting that there might be a mechanism to partially segregate the inner mitochondrial membrane without DRP-1 to maintain normal mitochondrial function to some degree. This may explain the lower level of *hsp-6p∷*GFP induction in *drp-1* mutant animals than in *fzo-1* mutant animals where mitochondria are completely fragmented and lost normal function. In *drp-1* mutant animals, ethanol cannot induce fragmentation of the outer mitochondrial membrane. However, considering that ethanol does further enhance *hsp-6p*∷GFP expression in *drp-1* mutant animals, it is possible that ethanol exposure may target the inner mitochondrial membrane for fission in the absence of DRP-1.

We showed that perpetual UPR^mt^ activation mediated by constitutively active ATFS-1 protects from alcohol-induced muscle dysfunction. Paradoxically, our data also indicate that ethanol exposure induces UPR^mt^ through ATFS-1 activation. Then, why would muscle function of ethanol-exposed animals decline even though ethanol activated a protective UPR^mt^? One possibility is that while ethanol’s ability to induce UPR^mt^ via ATFS-1 is not robust enough to protect from alcohol-mediated muscle weakness, constitutive activation of ATFS-1 with a gain-of-function mutation is sufficiently robust for such protection. Alternatively, UPR^mt^ induction prior to ethanol exposure is necessary to prevent cellular damage.

A simple mechanistic explanation for the protection from alcohol-mediated muscle dysfunction in *atfs-1(et15gf)* animals is that these animals produce a high level of ATP due to increased mitochondrial biogenesis. However, published data showed that ATFS-1 does not increase ATP production (10). ATFS-1 represses the transcription of oxidative phosphorylation components and TCA cycle enzyme genes but increases the transcription of glycolysis enzyme genes, thus maintaining overall ATP production. Furthermore, ATFS-1 regulates the biosynthesis of certain lipid species (32, 33). In addition to this metabolic adaption, ATFS-1 promotes oxidative phosphorylation complex assembly and an increase of protective mitochondrial chaperones and proteases designed to improve the health of mitochondrial proteome (9, 34). Importantly, this global change is beneficial to certain types of stresses or infection (32, 33, 35–37), but is potentially harmful to others (38, 39). Intriguingly, skeletal muscles of chronic alcohol users were reported to exhibit a reduction of aerobic metabolism and an increase of anaerobic metabolism (40). This may be explained by our findings of ethanol-mediated activation of ATFS-1 and UPR^mt^, which lead to metabolic adaptation. Based on all this information, we speculate that a branch of UPR^mt^ induced by ATFS-1 activation is directly involved in the protection from alcohol-mediated muscle dysfunction. Interestingly, although we do not know that UPR^mt^ influences alcohol-induced tissue damage, several studies in mammals indeed showed that UPR^mt^ upregulation is protective from certain stress-induced organ damage (41–43). Taken together, we propose that mitochondrial dysfunction and consequent UPR^mt^ activation may underlie many pathologies of alcoholic myopathy and further understanding of UPR^mt^ in alcoholic myopathy may lead to a better therapeutic intervention.

## Acknowledgements

We thank Pamela Hoppe for providing *myo-3*∷gfp plasmid construct. Some strains were provided by the *Caenorhabditis* Genetics Center, which is funded by the National Institutes of Health Office of Research Infrastructure Programs (P40 OD010440). This work is partially supported by funding from the National Institutes of Health (R21 AA023940 and RO1GM125749).

## Materials and Methods

### Worm strains and maintenance

All *C. elegans* strains were cultured at 20 °C on NGM (nematode growth medium) plates seeded with *E. coli* OP50. The following strains were used in this study: N2, SJ4100 (zcIs13[hsp-6p∷GFP]), SJ4005 (*zcIs4[hsp-4p*∷GFP]), CL2070 (*dvIs70[hsp-16.2p*∷GFP + *rol-6 (su1006)*]), CL2166 (*dvIs19[gst-4p*∷GFP∷NLS]) (44, 45), SJ4197(*zcIs39[dve-1p*∷*dev-1*∷GFP], CF2124(*muIs139[dod-11p*∷RFP(NLS) + *rol-6(su1006)*]), the infection reporter strain AU133 (*agIs17[irg-1p*∷GFP + *myo-2p*∷mCherry]) (46), the infection reporter strain AY101 (*acIs101[irg-5p*∷GFP + *rol-6(su1006)*]) (47), the xenobiotic detoxification reporter strain CY573 (*bvIs5[cyp-35B1p*∷GFP + *gcy-7p*∷GFP]) (48), the hypertonic stress reporter strain VP198 (*kbIs5[gpdh-1p*∷GFP + *rol-6(su1006)*]) (49), the proteasome stress reporter strain GR2183 (*mgIs72[rpt-3p∷GFP* + *dyp-5*(+)] (50), the hypoxia reporter strain DMS640 (*nIs470 [cysl-2p*∷GFP + *myo-2p*∷mCherry] (51), *drp-1(tm1108), fzo-1(tm1133), mff-1(tm2955), mff-2(tm3041), fis-1(tm1867), fis-2(gk414), dct-1(tm376),pdr-1(gk448),pink-1(tm1799), ced-3(n717), ced-9(n2812);ced-3(n717), atfs-1(gk3094lf), atfs-1(et15gf)*. Transgenic strains were constructed using standard DNA microinjection methods. The following transgenes were generated in our lab: *cimIs46[myo-3*∷GFP], *cimIs42[twk-28p∷atp-9*∷GFP], *cimIs40[myo-3p∷tomm-20*(mts)∷FusionRed], *cimEx102[atfs-1p∷atfs-1*∷GFP]

### Ethanol treatment

As described previously (52), the NGM agar plates used for all of the ethanol treatment experiments were dried without lids for 25 min in a 60 °C incubator. This drying process ensures rapid ethanol absorption into agar, and this process usually reduces the agar to 85-90 % of its original volume. After the plates were cooled down to room temperature, OP50 *E. coli* were seeded at a center area of the plate and were allowed to grow overnight. Next day, a desired amount of ethanol (300 mM), which gives an internal ethanol concentration of approximately 30 mM (15, 53), was applied to the NGM plates without direct contact with bacteria. The plates were then sealed with parafilm and left at room temperature for equilibration at least for an hour. The desired number of animals at the L4 stage were picked on the plate, and cultured for 24 h in a 20 °C incubator.

### Burrowing assay

The burrowing assay was performed as described by Lesanpezeshki et al. with some modifications (14). Briefly, we placed fifteen age-matched (24 h post L4) naïve or ethanol-exposed animals at the bottom center of a 12 well plate. We then added 2 ml of 26% (w/w) Pluronic F-127 solution and waited for 5 min to transition from liquid to gel at room temperature. Two microliters of isoamylalcohol (1:200 dilution in 70% ethanol) were spotted on the top surface, and then twenty microliter of OP50 bacterial culture was added on the dried isoamylacohol spot. Animals that successively burrowed to the top and were swimming on the bacterial culture solution were counted every fifteen minute and removed. Triplicate assays for each genotype and treatment were concurrently performed for 90 min.

### Thrashing assay

Fifteen animals were placed in 1 ml of M9 on a 35 mm unseeded NGM plate with care to minimize the co-transfer of the OP-50 bacteria. Worms were let sit for 5 minutes to become acclimated to the liquid environment. A video was recorded for 30 seconds using Image Pro 10. By replaying the video at a slower frame rate, the number of thrashing from individual worms was counted manually. Worms that remained in the field view for the entire 30 seconds were counted and the identity of the worms was blinded to the counting person. It was counted as one thrash when the head or tail of the worm swings back.

### Measurement of fluorescent reporter expression

Worms were mounted with 6 mM levamisole in M9 on a 2 % agarose pad. Images were captured using 10 x objective of Zeiss microscope and Metamorph. Using Metamorph, the outline of the worms was manually drawn and integrated pixel intensity was obtained. The background intensity, which was obtained from a neighboring region, was subtracted from the intensity of the sample.

### Mitochondrial morphology analysis

Worms were mounted on 2 % agarose pad using 6 mM levamisole in M9 solution. Images were captured using 63× objective of Zeiss microscope. First, the categorical score scale was established and by comparing to the established standard images, each muscle cell was given the score.

### Statistical analysis

All graphing and statistical analysis were performed using Prism 7. The sample numbers represent biological replicates.

---

**Movie S1**. Wild-type (WT) and *atfs-1(et15gf)* mutant animals, which were cultured in the presence or absence (control) of ethanol, show swimming movements (“thrashing”) in a liquid medium.

**Fig. S1.**
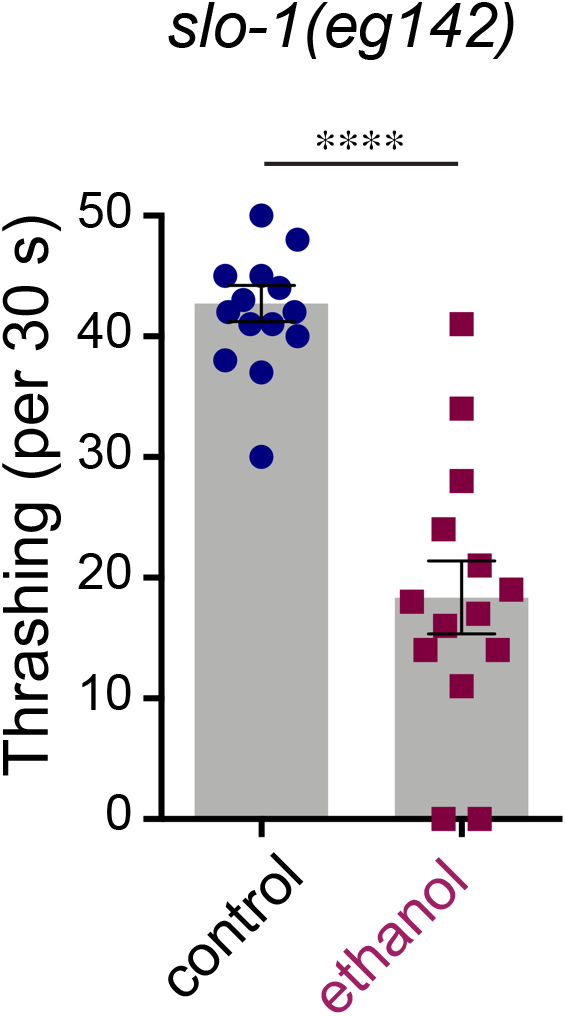
Animals resistant to the intoxicating effects of ethanol still show an impairment in thrashing. *slo-1(eg142)* mutant animals were exposed to ethanol for 24 hours. mean ± SEM ****p<0.0001, t-test

**Fig. S2.**
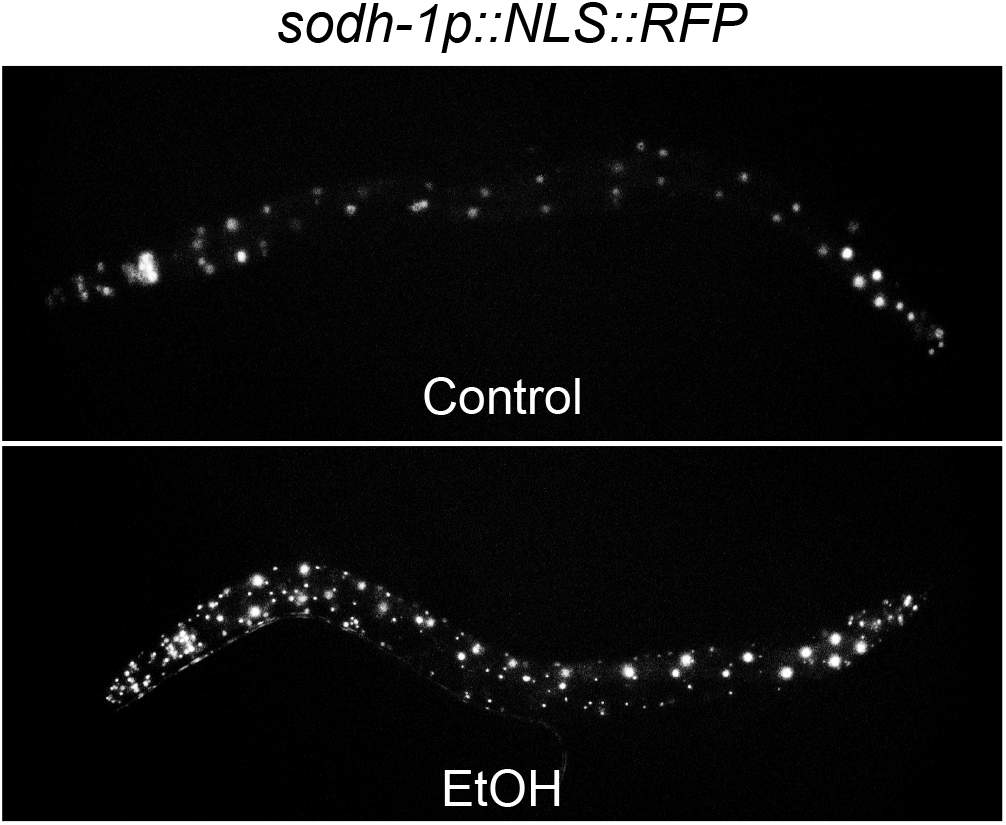
Ethanol strongly induces the expression of alcohol dehydrogenase *sodh-1*. Strain CF2124 (*muIs139 [dod-11p∷RFP(NLS)* + *rol-6(su1006)])* was exposed to ethanol from L4 stage for 24 hours. Scale bar:

**Fig. S3.**
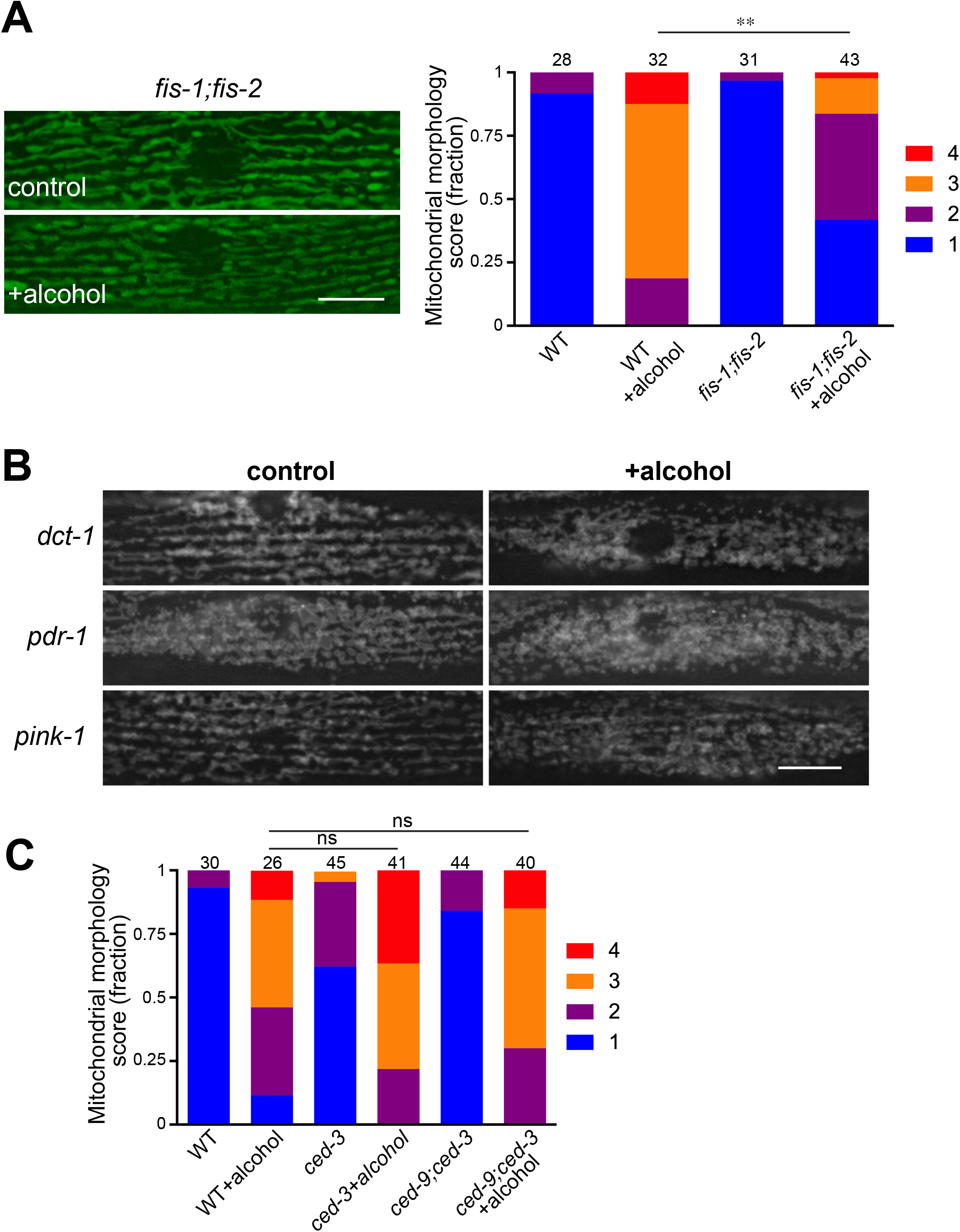
Mutations in genes that mediate mitophagy and cell death do not alter ethanol-induced mitochondrial fragmentation. *(A)* Mitochondrial networks in control and ethanol-exposed wild-type and *fis-1;fis-2* double mutant animals. Mitochondrial networks was assessed in the *cimIs42* background. Scale bar, 10 μm. **p < 0.01, Kruskal-Wallis test, Dunn’s post-hoc analysis. The numbers above the bar graph are the sample numbers. *(B)* Mitochondrial networks in control and ethanol-exposed *dct-1, pdr-1*, and *pink-1* mutant animals. These mutants have thick mitochondrial networks, which hinders accurate quantification. Scale bar, 10 μm. *(C)* Mitochondrial networks in control and ethanol-exposed wild-type, *ced-3, ced-9;ced-3* mutant animals. The numbers above the bar graph are the sample numbers.

